# Sequence and immunologic conservation of *Anaplasma marginale* OmpA within strains from Ghana as compared to the predominant OmpA variant

**DOI:** 10.1101/641696

**Authors:** JE Futse, G Buami, BB Kayang, R Koku, GH Palmer, T Graça, SM Noh

## Abstract

A primary challenge in developing effective vaccines against obligate, intracellular, bacterial tick-borne pathogens that establish persistent infection is the identification of antigens that cross protect against multiple strains. In the case of *Anaplasma marginale*, the most prevalent tick-borne pathogen of cattle found worldwide, OmpA is an adhesin and thus a promising vaccine candidate. We sequenced *ompA* from cattle throughout Ghana naturally infected with *A. marginale* in order to determine the degree of variation in this gene in an area of suspected high genetic diversity. We compared the Ghanaian sequences with those available from N. America, Mexico, Australia and Puerto Rico. When considering only amino acid changes, three unique Ghanaian OmpA variants were identified. In comparison, strains from all other geographic regions, except one, shared a single OmpA variant, Variant 1, which differed from the Ghanaian variants. Next, using recombinant OmpA based on Variant 1, we determined that amino acid differences in OmpA in Ghanaian cattle as compared to OmpA Variant 1 did not alter the binding capacity of antibody directed against OmpA Variant 1, supporting the value of OmpA as a highly conserved vaccine candidate.

## Introduction

The identification of protective antigens is a primary limitation of vaccine development against obligate, intracellular pathogens that use antigenic variation to establish persist infection within the mammalian host. These pathogens include, though are not limited to bacteria in the family *Anaplasmataceae*, such as *Anaplasma marginale, A. phagocytophilum* and *Ehrlichia ruminantium. A. marginale* causes bovine anaplasmosis which is a production limiting disease of cattle that occurs worldwide [1]. A single vaccine that protects against antigenically distinct variants is a high priority. This is particularly true for farmers in tropical regions of the world in which high infection prevalence and transmission pressure leads to pathogen diversification [2].

Whole outer membrane preparations induce protection against bovine anaplasmosis and in a proportion of vaccinates, protection against infection [3-5]. The immunodominant components of the outer membrane preparation include the major surface proteins Msp1, Msp2, Msp3, Msp4 and Msp5. Msp2 and Msp3 are best studied and are critical for immune escape and establishment and maintenance of persistent infection [6-8]. However, none of these major surface proteins have consistently produced protective immunity, though many trials have been conducted using both native and recombinant proteins [9-11].

Complete genome sequencing, proteomics and reverse vaccinology have identified a number of additional outer membrane proteins that could serve as vaccine candidates [12, 13]. However, narrowing the possible candidates is challenging in the absence of knowledge about proteins that mediate essential functions. Adhesins and invasins are particularly relevant vaccine candidates because they are functionally essential and tend to have highly conserved regions. Additionally, blocking adhesion and invasion to host cells is lethal for obligate intracellular pathogens.

Recently, *ompA*, annotated as Am854 in the *A. marginale* genome, was identified as an adhesin playing an essential role in *A. marginale* entry into mammalian and tick cells [14]. Consequently, this protein serves as a highly relevant vaccine candidate. In West Africa, where prevalence of *A. marginale* is up to 75%, there is a near complete absence of information concerning the extent of genetic variation of *A. marginale* in general and *ompA*, specifically [15]. Additionally, genetic variation is generally higher in regions where pathogen prevalence and transmission pressure are high [2, 16]. Thus, the genetic variation in *ompA* in such regions is a relevant question for vaccine development. In this paper we determine the variation in OmpA in *A. marginale* strains infecting cattle in Ghana and test whether this variation impacts antibody binding.

## MATERALS AND METHODS

### Animals and Sampling

The cattle used in this study were treated with strict accordance of guidelines set by University of Ghana Institutional Animal Care and Use Committee and the Washington State University Animal Care and Use Committee. The protocol was approved by University of Ghana Institutional Animal Care and Use Committee, by the Noguchi Memorial Institute for Medical Research’s (NIACUC protocol number: 2015-01-5X) and the Washington State University Animal Care and Use Committee (ASAF #3686).

Blood for DNA extraction and *ompA* amplification and sequencing was from cattle in areas with heavy tick infestation in the three major vegetative zones in Ghana in which cattle are reared. Serum for ELISAs was either from animals naturally infected under closely monitored field conditions with wild-type strains of *A. marginale* housed at the University of Ghana Livestock and Poultry Research Centre, Accra, Ghana or experimentally infected with the St. Maries strain of *A. marginale* at the Animal Disease Research Center, Pullman, WA. In both groups of animals, pre-immune serum was collected prior to infection and immune serum was collected at 40-50 days post infection at the time of control of acute rickettsemia.

### DNA extraction and amplification of *ompA* from whole blood

Genomic DNA was extracted from the peripheral blood samples collected in EDTA using a Gentra Puregene Blood kit (Qiagen, Germantown, MD) as described by the manufacturer’s protocol. PCR to amplify *ompA* was done using the following oligonucleotides: 5’- ATG CTG CAT CGT TGG TTA GC-3’ and 5’-TTC AGG CGC GAC CAC TCC TG −3’, and PCR master mix (Roche, Basel, Switzerland). Cycling conditions were as follows: initial denaturation at 94°C for 5 min, annealing at 60°C for 1 min, extension at 72°C for 1 min and final extension at 72°C for 7 min, for 35 cycles. The resulting PCR fragments were visualized by electrophoresis in 1% agarose gel stained with ethidium bromide.

All sequences have been deposited in GenBank and have accession numbers MK882853-MK882884.

### Cloning and sequencing of the recombinant OmpA

The amplicons were cloned into PCR 4 TOPO ^®^ Vector plasmids and TOP 10 *Esherichia coli* cells were transformed using the manufacturer’s protocol (Invitrogen, Carlsbad, CA). The resulting individual colonies were selected and grown overnight at 37°C using standard procedures. Plasmid DNA was extracted from the cells using the Q1Aprep ^®^ Spin Miniprep kit (Qiagen, Germantown, MD). To confirm the presence of the desired inserts, plasmid DNA samples were digested with EcoRI and the resulting fragments were visualized on a 1% agarose gel stained with ethidium bromide.

### Sequencing and analysis of *OmpA* genotypes

The plasmid DNA was sequenced using T3 and T7 primers and sequences were compiled and analyzed using the Vector NTI software package (Invitrogen, Carlsbad, CA). Sequences were aligned in multiples and pairwise fashions within and between the identified variants in Ghana and within and between sequences available in GenBank from *A. marginale* strains and isolates from N. America, Australia, Puerto Rico and Mexico [11], including the following: St. Maries (NC_004842.2), Puerto Rico (KM821238.1) [11], Dawn-Australia (AU) (KM821232.1) [17], Nayarit, Mexico (MX) N4506 (KM821237.1) and N3574 (KM821236.1) [2], Virginia (KM821238.1) [18], Kansas 6DE (KM821231.1) and EMΦ (KM821235.1) [19], and Colville-Washington C51 (KM821233.1) and C52 (KM821234.1) [11].

### OmpA Expression and Purification

Genomic DNA derived from the St. Maries strain of *A. marginale* was used as a template to amplify *ompA* as previously described [11]. The PCR product was cloned into pBAD202 Directional TOPO™ Expression vector (ThermoFisher Scientific, Waltham, MA) and inserted into TOP10 One Shot chemically competent cells. Positive transformation was confirmed by PCR and Sanger sequencing (Eurofins, Louisville, KY).

Protein expression, from a single randomly selected clone, was induced with 20% arabinose to a final concentration of 0.1% (v/v) once the OD600 reached 0.6 nm. The culture was grown for 4 hours, pelleted by centrifugation for 10 min at 10 000g and 4 °C. The pellets were flash frozen with liquid nitrogen and stored overnight at −80 °C. Pellets were resuspended in PBS containing cOmplete™ ULTRA Tablets, Mini, EDTA- free, Protease Inhibitor Cocktail (Roche, Basel, Switzerland) and sonicated for 4 min using a high intensity cup at 90% intensity (Fisherbrand™ Model 705 Sonic Dismembrator) in intervals of 20 seconds ON/OFF. The lysate was then centrifuged at 30,000 × g for 30 min and 4 °C.

The first step of purification was performed using 5mL of HisPur Nickel NTA resin (ThermoFisher Scientific, Waltham, MA) packed into a 10/10 GL Tricorn column (GE Healthcare, Chicago, IL) attached to an AKTA Avant HPLC apparatus (GE Healthcare, Chicago, IL). Sample loading was performed at 1mL/min and the remaining steps were performed at 2 mL/min. Chromatographic conditions were as follows: resin equilibration with 90% buffer A1 (50 mM Sodium phosphate 500 mM NaCl), 10% buffer B1 (50 mM Sodium phosphate 500 mM NaCl 150mM Imidazole pH 7.6) for 5 column volumes (CV); sample loading was performed using the sample pump; washout unbound for 5 CV in 10% Buffer B1, sample elution in 5 CV at 100% B1. Protein elution was detected measuring Abs at 280nm and fractions were collected in 1ml aliquots at 12°C. Pooled fractions correspondent to the entire peak that eluted at 100% B1 were buffer exchanged into Buffer A2 (20mM BisTris pH 6.5) using a Hi-trap G25 column attached to the AKTA and then chromatographed on a HPLC AKTA apparatus using a MonoQ GL 5/10 column (GE Healthcare, Chicago, IL). The Anion Exchange separation was performed at 1 mL/min with a using of Buffers A2 and B2 (20mM BisTris 250mM NaCl pH 6.5) and protein elution was detected by measuring Abs at 280 nm. Chromatographic conditions were: Isocratic (100% A2) for 5 CV, followed by sample loading using the sample pump, isocratic (100% A2) for 5 CV, 0-100% B2 for 25 CV, isocratic (100% B2) for 5 CV and isocratic (100% A2) for 10 CV. Protein peaks were collected using peak elution and protein purity and identity confirmed by SDS-PAGE and mass spectrometry (WSU central core facilities). Fractions containing pure OmpA were pooled and buffer exchanged by gel filtration using a G25 prepacked column attached to the AKTA into PBS pH 7.4. The data were collected, and the sample chromatograms were acquired using a Unicorn Chromatography Management System Version 7.0 (GE Healthcare, Chicago, IL). Sample concentration was determined by BCA (ThermoFisher Scientific, Waltham, MA). Successful protein purification was verified using standard SDS-PAGE (Fig.1).

**Fig. 1.**
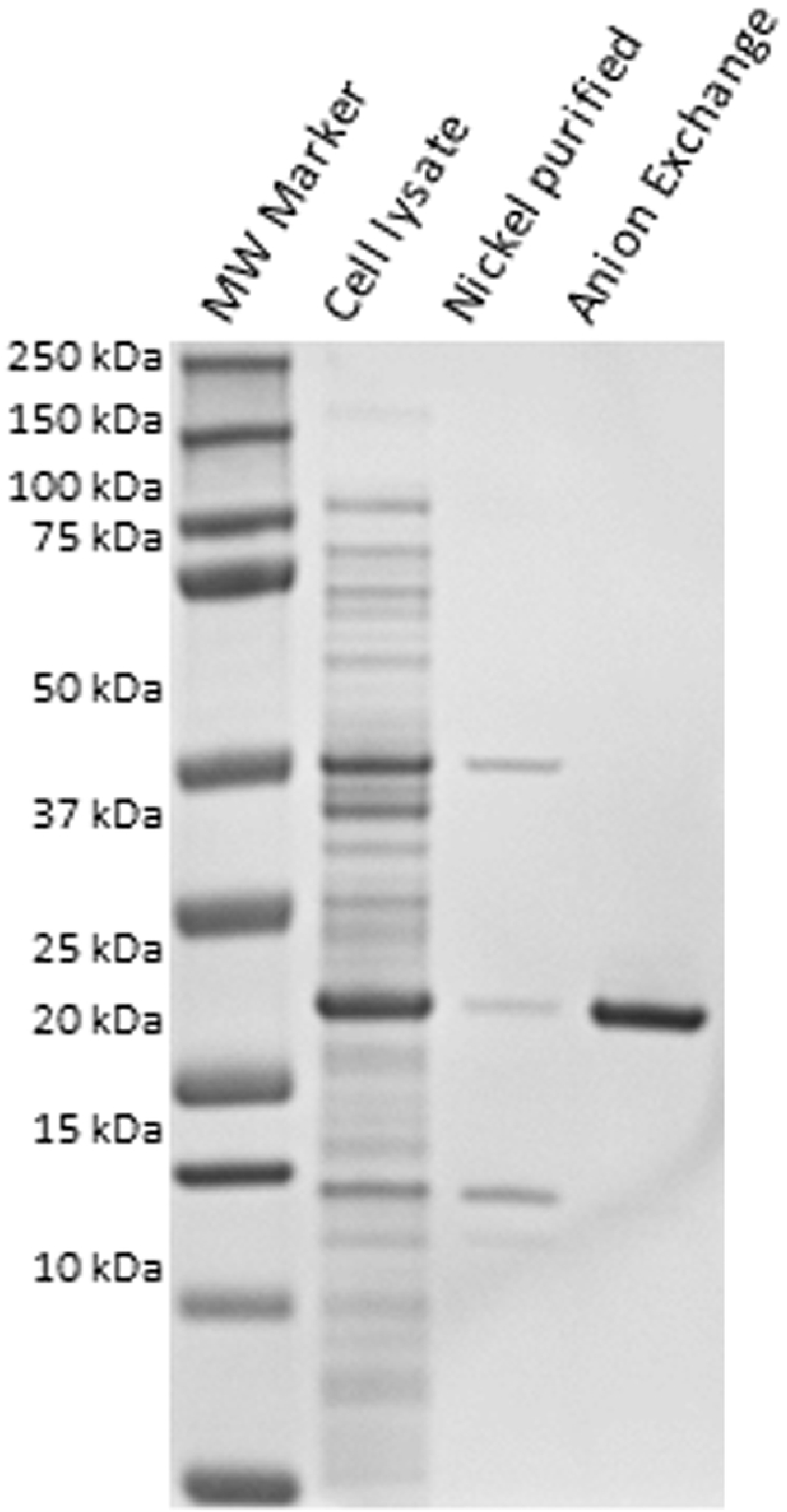
Expression and purification of recombinant OmpA. SDS-PAGE gel with recombinant OmpA prior to purification (cell lysate), following nickel column purification (nickel purified), and anion exchange chromatography (anion exchange). The expected molecular weight of OmpA is 26kDa.

### Enzyme-Linked Immunosorbent Assay (ELISA)

ELISA’s to detect anti-OmpA antibody were done as previously described [11]. Briefly, serial two-fold dilutions starting at 1:10 were used in triplicate to determine end-point titers. A positive titer was calculated as three standard deviations above the mean of the optical density of the pre-immune sera at the same dilution as the test sera [20]. The commercially available cELISA (VMRD, Pullman, WA) was used to measure anti-Msp5 antibody response using the manufacturer’s instructions.

### Statistical Analyses

All data analyses were conducted using GraphPad Prism (version 8.1.0 for Windows, GraphPad Software, La Jolla CA). End point titers for OmpA were log_10_ transformed. The Mann-Whitney two-tailed test was used to determine P values when comparing end-point titers directed and OmpA and percent inhibition of anti-Msp5 antibodies. A Mann-Whitney test was used to test null hypothesis that there is no greater diversity among Ghana strains than among the North America strains of *ompA*.

## RESULTS

### Conservation of *ompA*

Three new *ompA* variants were identified in *A. marginale* strains from cattle throughout Ghana (Table1). The overall conservation among the three variants was high with >99% nucleotide and >98% amino acid identity. None of the three variants were identical to the reference St. Maries strain or other previously reported *ompA* variants (Table 1, S1, S2) [11]. All Ghanaian variants have a synonymous substitution of a cytosine to a thymine at base pair 363, which is not found in other variants. Additionally, all Ghanaian variants have two substitutions that are shared only by Kansas EMΦ (Variant 3). The first of these is a synonymous substitution of guanine to adenine at base pair 663. The second is a non-synonymous substitution of thymine to cytosine at base pair 685, which results in an amino acid substitution of serine to proline. Ghana variant 1 (GV1) has only these three substitutions. Ghana variant 2 (GV2) has two additional non-synonymous substitutions. The first is a cytosine to thymine substitution at base pair 335 resulting in an amino acid substitution from a serine to a leucine. The second is at base pair 556 and consists of an adenine to guanine, resulting in a serine to glycine substitution. Ghana variant 3 (GV3) has one additional substitution of guanine to adenine at base pair 706 resulting in a glutamic acid to lysine substitution. Importantly the variation among the Ghanaian variants is no greater than among the other known variants (p >0.99).

**Table 1.** Location of codons with SNPs and the corresponding encoded amino acid in *ompA* variants from Ghana, N. America, Mexico, Puerto Rico and Australia.

| Variant | 335<sup>e</sup> | 112<sup>f</sup> | 363 | 121 | 531 | 177 | 556 | 185 | 663 | 221 | 685 | 229 | 706 | 236 |
| --- | --- | --- | --- | --- | --- | --- | --- | --- | --- | --- | --- | --- | --- | --- |
| GV1<sup>a</sup> | **TCG** | Ser | **TCT** | Ser | **GCG** | Ala | **AGC** | Ser | **GCA** | Ala | **CCG** | **Pro** | **GAA** | Glu |
| GV2 | **TTG** | **Leu** | **TCT** | Ser | **GCG** | Ala | **GGC** | **Gly** | **GCA** | Ala | **CCG** | **Pro** | **GAA** | Glu |
| GV3 | **TCG** | Ser | **TCT** | Ser | **GCG** | Ala | **AGC** | Ser | **GCA** | Ala | **CCG** | **Pro** | **AAA** | **Lys** |
| V1<sup>b</sup> | **TCG** | Ser | **TCC** | Ser | **GCG** | Ala | **AGC** | Ser | **GCG** | Ala | **TCG** | Ser | **GAA** | Glu |
| V2<sup>c</sup> | **TCG** | Ser | **TCC** | Ser | **GCA** | Ala | **AGC** | Ser | **GCG** | Ala | **TCG** | Ser | **GAA** | Glu |
| V3<sup>d</sup> | **TCG** | Ser | **TCC** | Ser | **GCG** | Ala | **AGC** | Ser | **GCA** | Ala | **CCG** | **Pro** | **GAA** | Glu |

<sup>a</sup>. GV1, GV2, GV3 represent the three variants from Ghanaian strains of *A. marginale*.
<sup>b</sup>. V1 represents St. Maries, Virginia, Kansas 6DE, Colville C51 and C52, and Nayarit (MX) N3574.
<sup>c</sup>. V2 represents Puerto Rico, Dawn (AU), Nayarit (MX) N4506.
<sup>d</sup>. V3 represents Kansas EMΦ.
<sup>e</sup>. Nucleotide position of the SNP within *ompA* as indicated by the bolded nucleotides within the codon.
<sup>f</sup>. Amino acid position encoded by the SNP. Bolded amino acids represent changes due to non-synonymous SNPs using V1 as the reference sequence.

There are three variants among all other previously identified *ompA* sequences. Variant 1 (V1) is the most common and is typified by the St. Maries reference strain. Variant 2 (V2) has one synonymous nucleotide substitution of a guanine to adenine at base pair 531, thus making the amino acid sequence identical to that of V1. V3, which represents only EMΦ, shares a single synonymous and a single non-synonymous SNP with all Ghanaian variants, as described above.

### Antibody Responses

In order to determine if these sequences differences affected antibody binding, OmpA V1, the most common variant, from the St. Maries strain of *A. marginale* was expressed as recombinant protein. The endpoint titers of the specific anti-OmpA response in animal infected with the St. Maries strain and the Ghanaian strains of *A. marginale* were compared. Sequence differences between OmpA V1 and the Ghanaian strains did not alter antibody binding (Fig. 2A). In animals from Ghana, titers to OmpA V1 tended to be higher than in animals infected with the homologous St. Maries strain, though there was marked variation between animals in both groups. All animals in Ghana had titers to OmpA. The titers varied from 10 to 12,800 with a mean titer (+/− CI) of 2459 +/− 3437. In the St. Maries-infected animals, three animals lacked titers to OmpA. The maximum titer was 640. The mean titer (+/− CI), including the negative animals was 166 +/−149.30.

**Fig 2.**
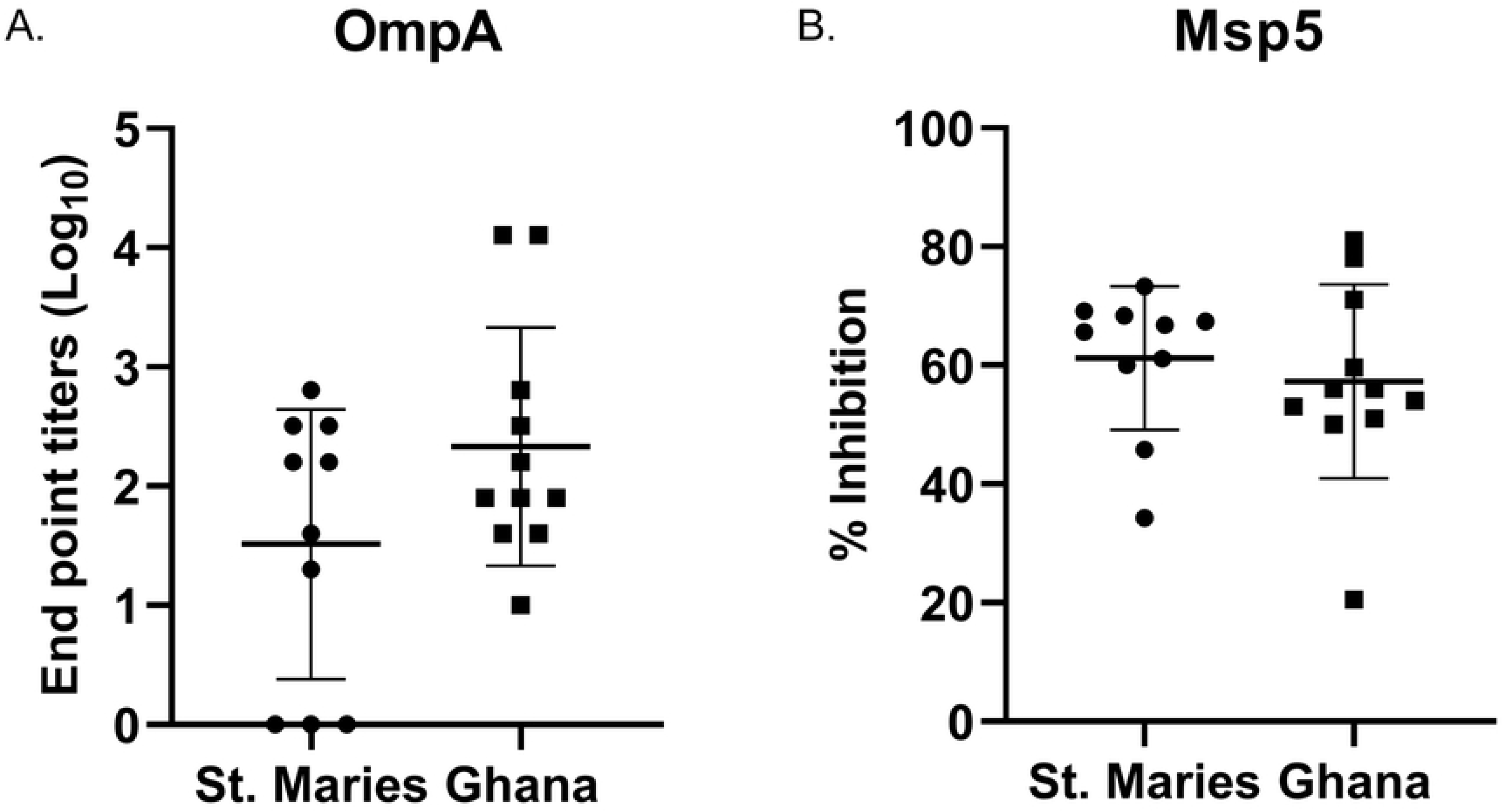
Magnitude of the antibody response directed against *A. marginale* OmpA V1 and Msp5. A. Anti-OmpA antibody response directed against recombinant OmpA V1 in animals infected with the homologous St. Maries strain or Ghanaian strains. B. Anti-Msp5 antibody response in animals infected with the St. Maries strain or Ghanaian strains of *A. marginale*. The horizontal bars represent the means of each population. The vertical lines represent the standard deviation.

A commercially available cELISA was done to verify that the overall higher titers in the Ghanaian animals did not reflect generally elevated titers to all *A. marginale* proteins (Fig. 2B). This cELISA detects antibody against the highly conserved Major Surface Protein 5. The percent inhibition in animals from Ghana was between 21-81% (mean +/− CI = 57% +/−10.96). The percent inhibition in animals infected with the St. Maries strain was between 34 and 73% (mean +/− CI= 61% +/−8.64).

## Discussion

One of the major limitations of developing a vaccine against *A. marginale* is strain diversity and the inability to produce cross-protection against a variety of field strains with a single vaccine. Consequently, vaccine development and antigen selection have focused on the identification of outer membrane proteins that are broadly conserved across distant geographic areas. For example, Omp 5, 9, 12, 13 and 14 are invariant during acute and persistent infection as well as tick transmission [21]. However, all of these Omps have variation between strains even when comparing strains from N. America including St. Maries, Florida, S. Idaho, Washington-Okanogan, and Oklahoma [22]. Specifically, for Omp5, the maximal number of nucleotide substitutions is 46 when St. Maries and FL strains were compared. Similarly, there are 44 substitutions in Omp9 when comparing the St. Maries and S. Idaho strains. In Omp14, there are 38 substitutions between Washington-Okanogan and S. Idaho. There are 17 substitutions in Omp13 when comparing the S. Idaho and Florida strains and 11 substitutions in Omp12 when comparing Oklahoma and S. Idaho. Consequently, this variation potentially limits these Omps as vaccine candidates.

Interestingly, this study demonstrates that OmpA has less variation than these other outer membrane proteins, with a maximum of three amino acid differences between the most common OmpA variant, V1, and the Ghanaian variants. Additionally, these amino acid differences did not alter the capacity of antibody raised against the Ghanaian strains to bind the heterologous OmpA V1 supporting the value of OmpA as a highly conserved vaccine candidate. Importantly, the endpoint titers are widely variable between animals. For example, in animals infected with Ghanaian strains, titers vary from 10 to 12,800, and in St. Maries infected animals titers vary from 0 to 640. This difference reflects the expected variability in the immune response in outbred animals, which must be addressed in vaccine development.

The reason for the overall higher titers directed against recombinant OmpA in the Ghanaian animals is unknown. However, to ensure that this difference does not reflect anti-*A. marginale* antibodies levels in general, the commercially available cELISA was used to evaluate the anti-msp5 antibody response. Msp5 is an immunodominant, highly conserved outer membrane protein, commonly used for diagnostic purposes [23]. The % inhibition between groups of animals was similar with mean of 61% and 57% for the St. Maries- strain infected and Ghanaian strain –infected animals, respectively.

OmpA serves as an adhesion and invasion for *A. marginale* and as such is an important vaccine candidate [14]. Here we demonstrate that OmpA is remarkably conserved in *A. marginale* from geographically diverse regions and the minimal difference between OmpA sequences does not affect antibody binding. Additionally, no amino acid diversity is present in the experimentally determined binding domain [14]. In one study, recombinant OmpA failed to induce protective immunity against tick challenge in cattle[11]. However, the binding domain is not a predicted B cell epitope and, in that study, the overall antibody response to a peptide containing the binding domain was low. Thus, appropriate adjuvants and delivery strategies need to be developed to overcome the marked variation in the immune response between animals and to maximize the potential protective capacity of the anti-OmpA immune response.

## Acknowledgements

This work was supported by the Wellcome Trust for the fellowship in Public Health and Tropical Medicine grant 097171/Z/11/Z, NIH Grant (R37 AI44005), USDA-ARS 2090- 32000-038-00D, and the Jan and Jack Creighton Endowment fund. We gratefully acknowledge the technical help of Jonathan Quaye, Deb Alperin and Jessie Ujczo.

